# Visual and Instrumental Assessment of Interaction of UVC Radiation with Liposomes in FeCl_3_ Solutions

**DOI:** 10.64898/2026.08.21.746308

**Authors:** Vladimir M. Subbotin, Benjamin A. Turner, Brian A. Davies, Kevin Wu, Gennady Fiksel

**Affiliations:** Arrowhead Pharmaceuticals Inc., Madison, WI 53719, USA; University of Wisconsin, Department of Human Oncology, Madison, WI 53792, USA; University of Wisconsin, Department of Physics, Madison, WI 53706, USA

**Keywords:** origin of life, liposomes, solar UV

## Abstract

Previously, we have demonstrated that certain ferric salts common in Archean waters, such as iron trichloride and ferric ammonium citrate, can protect liposomes from destruction by short-wavelength UVC light. In this study, we investigate the propagation of 254 nm UV radiation through aqueous FeCl_3_ solutions and its interactions with liposomes. We then consider these findings in the context of early Earth’s UV environment, discuss their implications for our hypothesis of the Darwinian evolution of liposomes, and integrate them with our previous experimental results.

## 1. Introduction

The self-assembly of bilayer lipid membranes into liposomes on the surface of Archean ponds provided the initial vital compartmentalization events needed for the origin of life, serving as protective physical boundaries and concentrating early organic molecules, such as phosphorylated ribose, nucleosides and nucleotides [1–5]. However, the harsh UV environment of the early Earth [6] can damage and destroy the surface-located liposomes unless, by encapsulating heavy solutes, they sub-merge and escape UV damage. As a result, through the processes of congregation, fusion, fission, and exchange of aggregated material, primitive Darwinian evolution can be enabled [7]. Thus, cyclic solar UV exposure combined with gravity can also be an environmental driver in the prebiotic evolution of liposomes [7,8].

It is known that UV absorption in pure water is very low [9,10], and that it requires certain impurities to provide sufficient shielding. Therefore, we investigated [10] the attenuating ability of several ferric salts typical for Archean waters. We showed that aqueous solutions of iron trichloride and ferric ammonium citrate, at concentrations thought to be present in Archean waters [11] attenuate UV intensity by a factor of 1000 at submersion depths of a few millimeters.

In a follow-up study [8], we used specially designed UV-sensitive liposomes (Encapsula NanoSciences, Brentwood, USA) to demonstrate that the UVA part of the UV spectrum does not destroy UV-sensitive liposomes, while exposure to UVC radiation results in liposome destruction.

The objective of this study is to further quantify the propagation of UV in ferric salt FeCl_3_ solutions and visualize and quantify the UV interaction with UV-sensitive liposomes. In particular, we explore several scenarios for liposome submersion, ranging from surface localization to complete submersion, to uniform dispersion throughout.

## 2. Material and Methods

### 2.1. Liposomes

We used large polymerizable UV-sensitive multilamellar micron-sized liposomes prepared by Encapsula NanoSciences, Brentwood, TN, USA, and composed of Dibehenoyl-sn-glycero-3-phosphocholine (DBPC) and 10,12-pentacosadiynoic acid (PCDA) with a DBPC: PCDA molar ratio of 80:20, at a total lipid concentration of 20 mM. The average liposome diameter was measured to be 1.6 μm.

The mechanisms of liposomal UV sensitivity and UV destruction were described in [8]. Briefly, the PCDA is a polymerizable unsaturated fatty acid. PCDA lipids adjacent to each other polymerize upon UV radiation, changing their color from the original white to blue [12]. According to the liposome manufacturer, the dynamics of the liposome color change is a reliable indicator of liposome destruction. The color change was assessed visually and imaged using a KEYENCE BZ-X810 microscope at x400 magnification. It was also quantified by measuring the absorbance of the emitted light from a BioTek Synergy Neo2 Hybrid Multimode Reader (Agilent, Santa Clara, CA, USA). The absorbance data were acquired by a PerkinElmer Synergy Neo2 plate reader.

### 2.2. UV Source

We used a mercury low-pressure 15-Watt Spectroline Germicidal UV Sterilizing lamp (General Tools, USA) with an intensity in the UVC range (254 nm) of 9.5 mW/cm^2^ at a distance of 30 mm from the lamp as measured by a UV512C Light Meter (General Tools, USA). It was demonstrated [8] that the UVA part of the UV spectrum does not destroy UV-sensitive liposomes, while the UVC part, in particular 254 nm emitted by a low-pressure mercury lamp, results in liposome destruction. The destruction was visually and instrumentally quantified by the liposomes’ color change due to their membranes’ interaction with light.

### 2.3. UV Shielding Medium

Based on our previous observations, we used a water solution of iron trichloride FeCl_3_ at a concentration varying from zero to 2.5 g/L.

### 2.4. Measurement of UV Absorption Spectrum

The wavelength dependence of the UV absorption in FeCl_3_ solutions was measured by a multicell Peltier Module G9889A for the Agilent Cary 3500 UV-Vis Spectrophotometer.

### 2.5. Liposome Dispersion

To study UV interaction with liposomes, we used a 96-well plate (Greiner, Austria) filled up to a depth of 10 mm with a pre-measured FeCl_3_ solution, after which 30 μL of undiluted liposomes were added. Three scenarios of liposome dispersion were studied. In the first scenario, the liposomes were layered onto the solution surface 14 hours prior to UV irradiation. We called it the night-time scenario because it simulates liposome formation at night, giving them sufficient time to submerge. In the second, so-called daytime scenario, the liposomes were slowly added during UV irradiation, which lasted typically from three to six minutes. Finally, a mixing scenario was accomplished by carefully dispersing the added liposomes to achieve a uniform distribution throughout the solution prior to UV irradiation. In nature, this scenario could be realized by large-scale water motion driven by wind or by soil disturbances.

To achieve uniform distribution of liposomes, we employed a specialized design to enable slow liposome delivery, which is especially important under UV radiation. The design comprises three 5-cm-long 33-gauge hypodermic tubing segments, extended to the centers of the targeted wells, with tips curved so that they touch the solution surface - Fig. 1. We used three 50 μL Hamilton syringes and a Harvard Apparatus syringe pump to deliver liposomes slowly during UV irradiation. This design ensures a smooth entry of the liposome into the solution, thereby eliminating the effect of initial liposome velocity on the submergence, which is then governed solely by gravity, buoyancy, and the solution viscosity.

**Figure 1.**
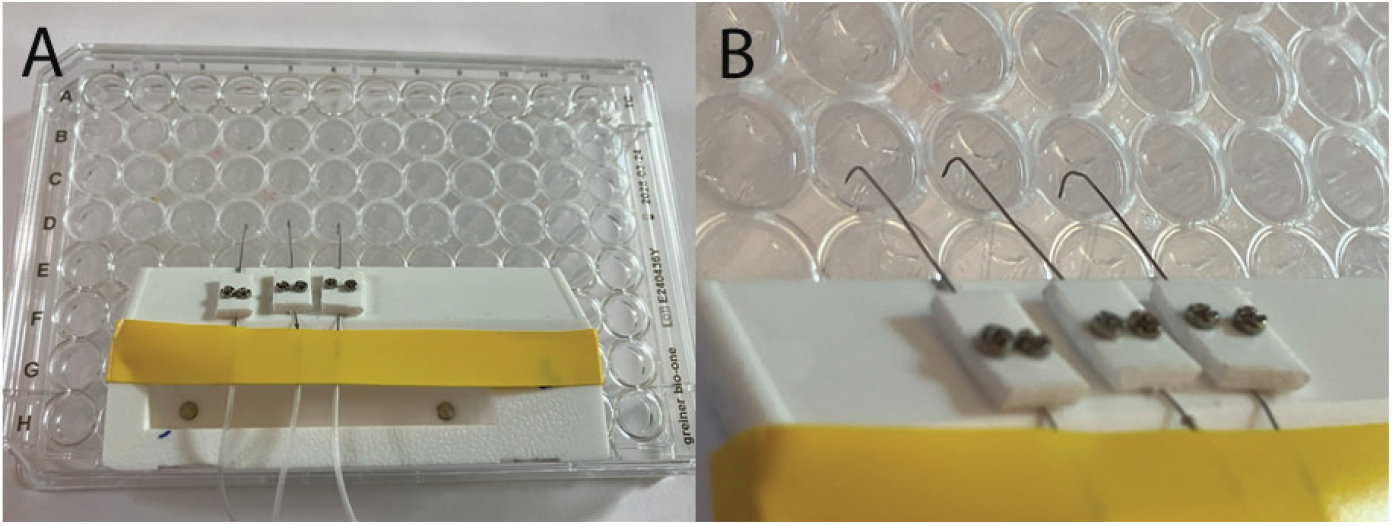
A device for slow delivery of liposomes during UV radiation (daytime scenario). A - general view,B-a magnified portion showing the curved tips.

## 3. Results

### 3.1. UV Absorption Spectrum

A UV absorption spectrum was measured at a FeCl_3_ concentration of 0.10 g/L and a UV light pass distance of *s* = 4 mm. A plot of absorption *A* = log_10_ *I*_0_/*I* vs the UV wavelength, where *I*_0_ and *I* are respectively the intensities of the incident and passing light, is shown in Fig. 2. The absorption value at a wavelength of 254 nm is *A* = 0.78. Assuming the absorption linearly depends on the solute concentration *ρ* and the light pass distance *d, A* = *ϵρd*, the absorption coefficient *ϵ* = *A*/*ρd* = 1.95 L/g mm. Conversely, a distance *d* at which the absorption reaches a certain value *A* can be found from a known absorption coefficient *ϵ* and the solute concentration *ρ*, so *d* = *A*/*ϵρ*.

**Figure 2.**
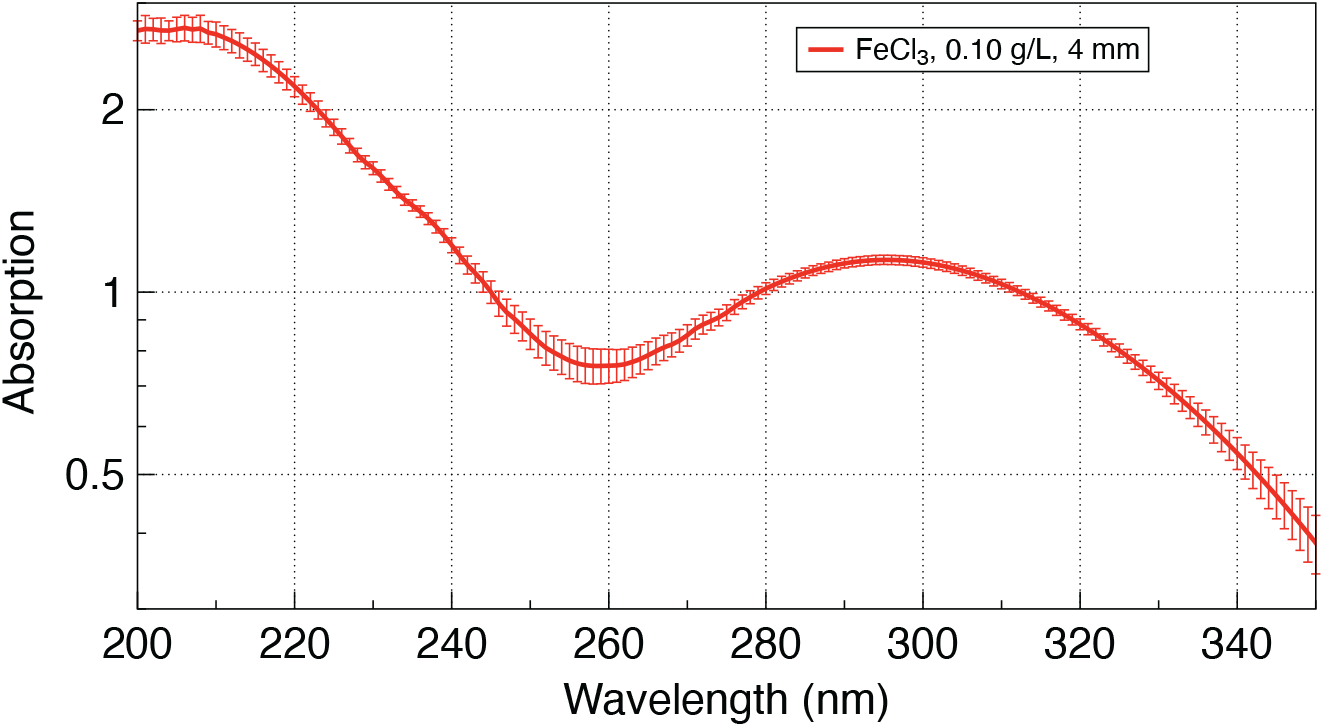
Absorption of UV light in 4 mm of FeCl_3_ solution at a concentration of 0.10 g/L. The error bars indicate a standard deviation over three consecutive measurements.

### 3.2. Night-Time and Daytime Scenarios

A qualitative visualization of the UV damage during the night-time and daytime scenarios is illustrated in Fig. 3. As mentioned before, the damage is associated with the intensity of the liposomes’ blue color after UV exposure. The results showed that liposomes added to the wells 14 hours prior to UV radiation to the wells filled with pure water were destroyed entirely (wells D 10-12) as indicated by the intense blue tint, while liposomes added 14 hours prior to UV radiation to the wells with iron trichloride solution so they would have sufficient time to submerge were completely protected (wells D 7-9). Liposomes that were added to the wells with iron trichloride solution during UV radiation do not have sufficient time to submerge and remain localized near the surface. In this case, they also sustained some damage, as indicated by a color change, although the extent was less than that of liposomes submerged into pure water.

**Figure 3.**
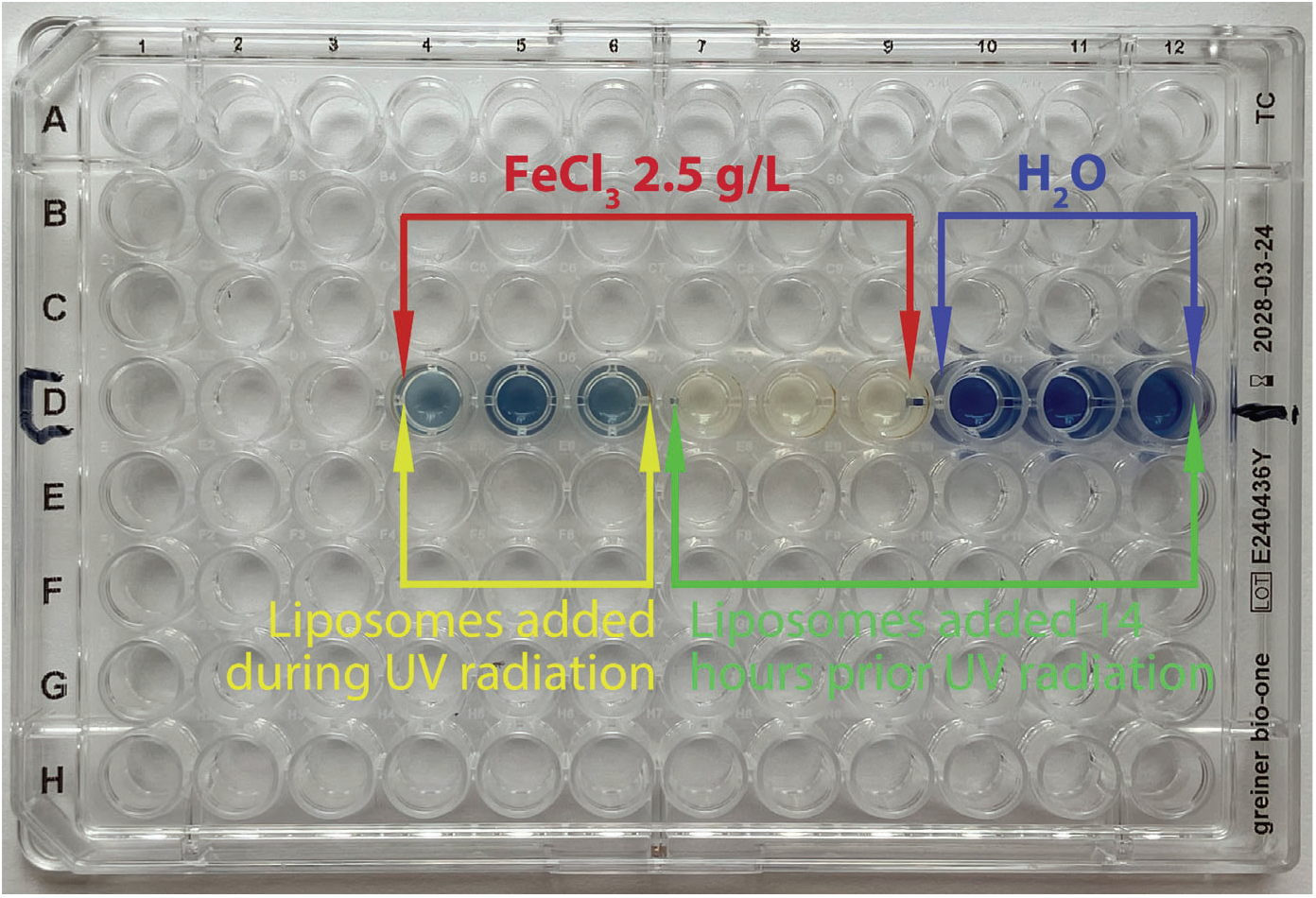
A photo of the experimental setup with a 96-well plate after 6 minutes of UV radiation. Liposomes submerged into pure water are completely destroyed (D 10, 11, 12), while liposomes shielded by FeCl_3_ solution appeared to be completely protected (D 7, 8, 9). Liposomes added to FeCl_3_ solution during UV radiation also appear to be destroyed (D 4, 5, 6), although by a lower amount, as evidenced by a lower degree of change in blur color intensity.

### 3.3. Mixing Scenario

In the mixing scenario, the liposomes were added to FeCl_3_ solutions and carefully dispersed to ensure uniform distribution throughout the solution prior to UV irradiation. In nature, this scenario could be realized by large-scale water motion driven by wind or by soil disturbances. For this series, we used the NUNC MaxiSorp 96 Well Plates (Thermo Fisher Scientific) with the outer front wall removed. The solution was exposed to a radiation intensity of 9.5 mW/cm^2^ for 3 min, which is equivalent to a UV dose of 1.7 J/cm^2^. A side view photo of the wells with the lines indicating calculated positions of absorptions A = 1 (1/10 attenuation), A = 2 (1/100 attenuation), and A = 3 (1/1000 attenuation) is shown in Fig. 4. The FeCl_3_ concentration is shown above each well. The images indicate that liposome damage gradually decreases with depth and then disappears at a UV absorption depth between that of *A* = 2 and *A* = 2. Table 1 gives corresponding calculated depths at different FeCl_3_ concentrations.

**Table 1.** The submersion depth *d* [mm] corresponding to different absorption *A* values.

| FeCl<sub>3</sub> concentration (g/L) | $d(A = 1)$ | $d(A = 2)$ | $d(A = 3)$ |
| --- | --- | --- | --- |
| 0.15 | 3.4 | 6.8 | 10.2 |
| 0.30 | 1.7 | 3.4 | 5.1 |
| 0.60 | 0.85 | 1.7 | 2.6 |
| 1.25 | 0.41 | 0.82 | 1.2 |
| 2.5 | 0.2 | 0.4 | 0.6 |

**Figure 4.**
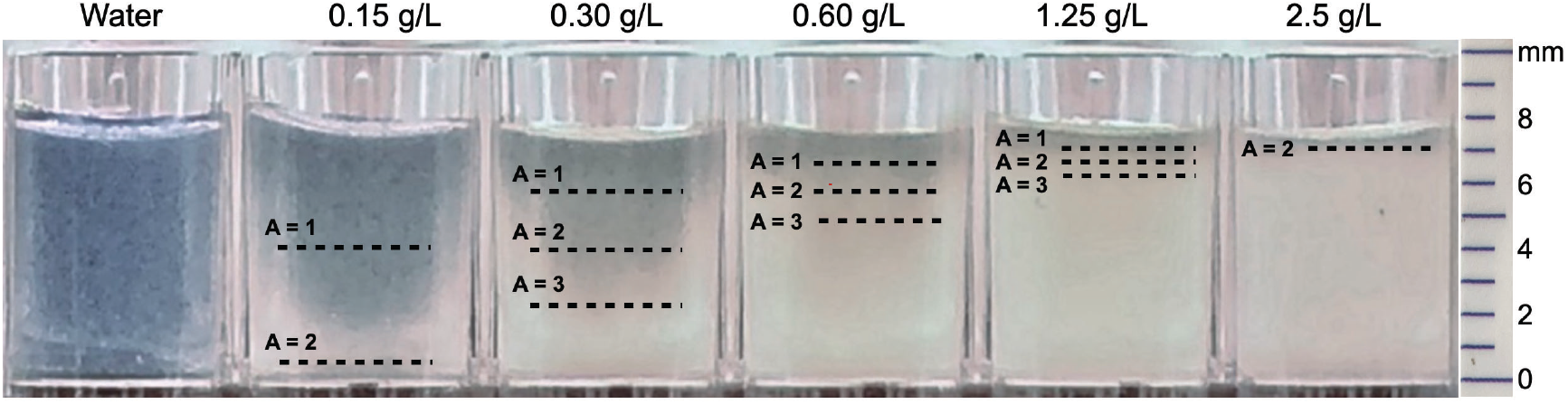
Side view photo of the wells with different FeCl_3_ concentrations. The dashed lines indicate calculated positions of absorptions *A* = 1 (1/10 attenuation), *A* = 2 (1/100 attenuation), and *A* = 3 (1/1000 attenuation).

### 3.4. Instrumental Assessment of Liposome Destruction by UV

The described visual evaluation of liposome destruction by UV was complemented by analyzing the light absoption technique using a BioTek Synergy Neo2 Hybrid Multimode Reader, as previously described [8]. First, the UV-irradiated samples shown in Fig. 4 were carefully mixed to obtain a uniform distribution of the blue-colored media as shown in Fig. 5.

**Figure 5.**
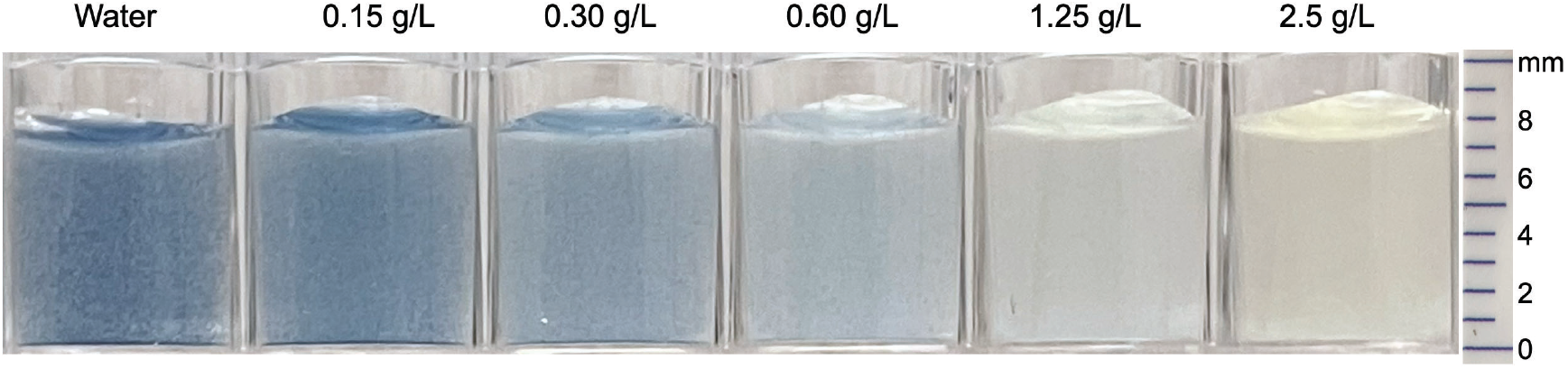
Mixed Post-UV samples

Afterward, the samples were transferred to the BioTek Synergy Neo2 Hybrid Multimode Reader for spectral measurements. The preliminary mixing and evaluation of absorption across the entire sample depth provide an estimate of the total number of destroyed liposomes. An example of several absorption spectra over a range of 450 nm to 750 nm is shown in Fig. 6. The absorption in pure water is very low. When liposomes are added, the absorption increases across the whole spectral range due to the light scattering on the liposomes. When FeCl_3_ is added, the absorption also increases somewhat due to a change in the solution color. However, no absorption peaks are observed before the UV exposure.

**Figure 6.**
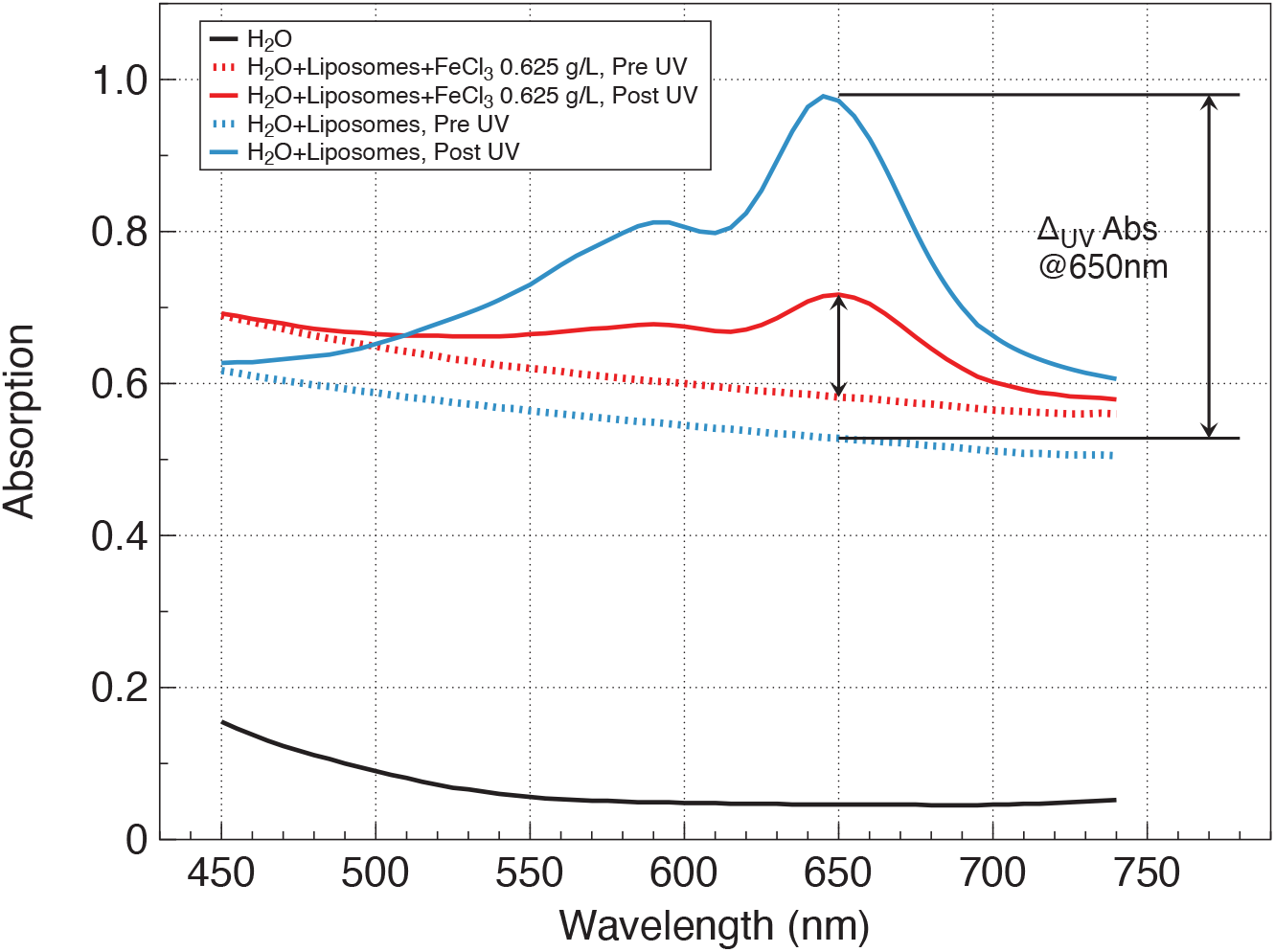
Absorption spectra for 3 cases: 1) Pure water, 2) Water with dispensed liposomes, and 3) Water with dispensed liposomes and added FeCl_3_. Dotted lines indicate pre-UV measurements and solid lines are the result of UV exposure. The incremental increase of absorption at 650 nm from pre-UV to post-UV, Δ_*UV*_ Abs is indicated by the vertical arrows.

After UV exposure, the solution becomes colored blue due to the damage to some liposomes, and the absorption spectra show a peak around 650 nm as well as a smaller peak at 595 nm resulting from preferred absorption of the red-orange part of the spectrum in a blue-colored medium. As a measure of the UV-induced damage, we take the incremental increase of absorption at 650 nm from pre-UV to post-UV, as indicated by the vertical arrows in Fig. 6. Its dependence on FeCl_3_ concentration is shown in Fig. 7. The result indicates progressively increased UV absorption, thus reduced UV damage, with increasing FeCl_3_ concentration.

**Figure 7.**
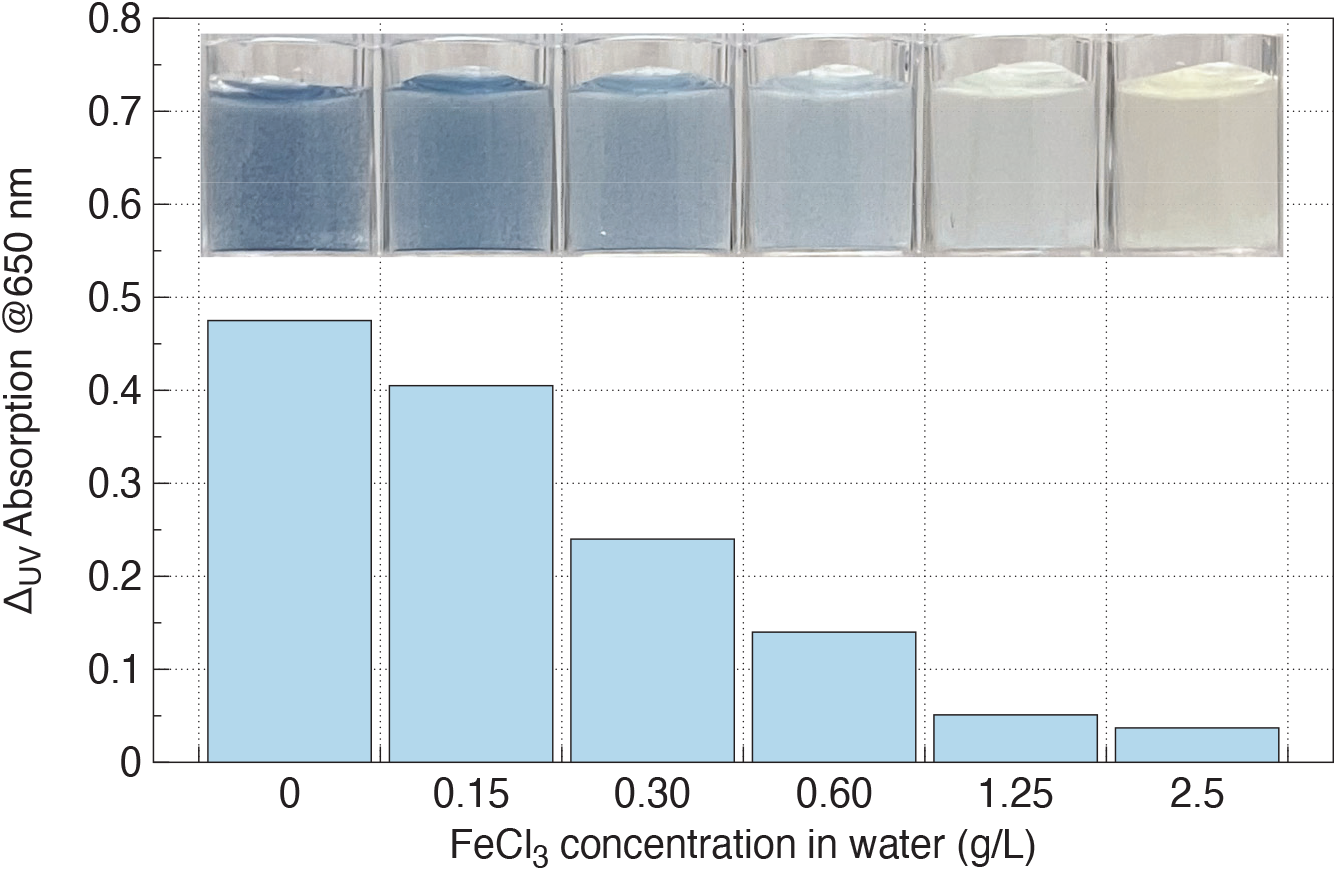
Incremental increase of absorption at 650 nm, Δ_*UV*_ Abs, from pre-UV to post-UV vs FeCl_3_ concentration.

## 4. Conclusions and Discussion

### 4.1. Conclusions

In this study, we have visualized and analyzed several scenarios of liposome interaction with UVC radiation. Three scenarios of liposome mixing-in were studied. In the first scenario, the liposomes were layered onto the surface of the iron trichloride solution 14 hours prior to UV irradiation. We called it the night-time scenario, simulating liposome formation at night and allowing sufficient time to submerge. In the second, so-called daytime scenario, the liposomes were slowly added during UV irradiation, typically from three to six minutes. In this scenario, the liposomes do not have sufficient time to submerge and remain localized near the surface. Finally, a mixing scenario was accomplished by carefully dispersing the added liposomes to achieve a uniform distribution throughout the solution. In nature, this scenario could be realized by large-scale water motion driven by wind or by soil disturbances.

We used large polymerizable UV-sensitive multilamellar micron-sized liposomes prepared by Encapsula NanoSciences, Brentwood, TN, USA, and composed of Dibehenoyl-sn-glycero-3-phosphocholine (DBPC) and 10,12-pentacosadiynoic acid (PCDA) with a DBPC: PCDA molar ratio of 80:20, at a total lipid concentration of 7 mM or 20 mM. The mechanisms of liposomal UV sensitivity and UV destruction were described in [8]. Briefly, the PCDA is a polymerizable unsaturated fatty acid. PCDA lipids adjacent to each other polymerize upon UV radiation, changing their color from the original white to blue [12]. According to the liposome manufacturer, the dynamics of the liposome color change is a reliable indicator of liposome destruction.

The liposomes were irradiated by UVC light with a wavelength of 254 nm at an intensity of 9.5 mW/cm^2^ and a time duration of 3 min. In all scenarios, the liposome damage was sensitive to the FeCl_3_ concentration but even at the lowest concentrations used, submersion by a few millimeters was sufficient to avoid the UV damage.

To quantify the destruction of liposomes by UV, we complemented the visual assessment with analysis of absorption spectra of the irradiated species. The solution turns blue in color due to interaction of liposome membranes with UV, and that results in the appearance of the absorption peak in the red-orange spectral part of the scanning light. The peak reaches its maximum in pure water and decreases substantially as the FeCl_3_ concentration in the liposomal solution increases, signifying progressively reduced UV-induced damage.

### 4.2. Discussion

How do the obtained results relate to the environment of Archean Earth? In the lab, at an intensity of 9.5 mW/cm^2^ a complete destruction of UV-sensitive liposomes was reached at 3 min, which is equivalent to a UV dose of *D*_*lab*_ = 1.7 J/cm^2^. During the Archean period, according to [6], the intensity of UV light in the UVC range (200 – 280 nm) was 0.31 mW/cm^2^, corresponding to a total UV dose during a 6-hour daytime of *D*_*Earth*_ = 6.7 J/cm^2^, which is *D*_*Earth*_ /*D*_*lab*_ = 4 times higher than the lab dose. However, due to the exponential decay of UV radiation in FeCl_3_ solution, that dose increase can be negated by a corresponding increase in absorption, that is, an increase in the submersion depth.

A precise calculation of the required depth is complicated because it is a convolution of the actual UV spectrum of Archean Earth, UV absorption spectra, and its biological activity. Given that the absorption at 254 nm is near the local minimum of the absorption curve in Fig. 2, assume, as the worst-case scenario, the same wavelength-averaged absorption coefficient *ϵ* for environmental UV. The absorption increase can be estimated as Δ*A* = log_10_ *D*_*Earth*_ /*D*_*lab*_ = log_10_ 4 = 0.6. Thus, to attain protection comparable to lab submersion at *A*_*lab*_ = 3, for example, the liposomes on Archean Earth must be submerged to an absorption value of *A*_*Earth*_ = 3.6, representing a 20% increase. Therefore, even if in a hypothetical scenario where the daily UV radiation dose is ten times higher than in the lab, achieving protection equivalent to submersion at *A*_*lab*_ = 3 in the lab would require the liposomes to be submerged to a depth only 20% greater than the values listed in the last column of Table 1.

The above results lend further credence to the novel hypothesis of the Origin of Life and the Darwinian evolution of liposomes [7].

## Author Contributions

Conceptualization, V.S. and G.F.; methodology, V.S. and G.F.; investigation, V.S., B.D., B.T., K.W. and G.F. ; data curation, V.S., B.D., B.T., K.W. and G.F. ; writing – original draft preparation, V.S. and G.F.; writing – review and editing, V.S., B.D., B.T., K.W. and G.F. ; visualization, V.S., B.D., B.T., K.W. and G.F. ; All authors have read and agreed to the published version of the manuscript.

## Acknowledgments

The authors want to thank Dr. Zahra Mirafzali (Encapsula NanoSciences, Brentwood, TN) for valuable advice and technical support.

## Data Availability Statement

The data supporting the findings of this study are available from the corresponding author upon reasonable request.

